# Pro-inflammatory cytokines disrupt β-cell circadian clocks in diabetes

**DOI:** 10.1101/705210

**Authors:** Naureen Javeed, Matthew R. Brown, Kuntol Rakshit, Tracy Her, Zhenqing Ye, Jeong Heon Lee, Tamas Ordog, Aleksey V. Matveyenko

**Author notes:** Corresponding author: Aleksey V. Matveyenko, PhD, Department of Physiology and Biomedical Engineering, Mayo Clinic, 200 First Street SW, Rochester, MN, 55905.

## Abstract

Intrinsic β-cell circadian clocks are a prerequisite for the control of glucose homeostasis through regulation of β-cell function and turnover. However, little is known about the contributions of circadian clock disruption to the natural progression of β-cell failure in diabetes. To address this, we examined the effects of cytokine-mediated inflammation, common to the pathophysiology of Type 1 and Type 2 diabetes, on the physiological, molecular, and epigenetic regulation of circadian clocks in β-cells. Specifically, we provide evidence that the key diabetogenic cytokine IL-1β disrupts functionality of the β-cell circadian clock and circadian regulation of insulin secretion through impaired expression of the key transcription factor Bmal1, evident at the level of promoter activation, mRNA, and protein expression. Additionally, IL-1β-mediated inflammation was shown to augment genome-wide DNA-binding patterns of *Bmal1* (and its heterodimer, *Clock*) in β-cells towards binding sites in the proximity of genes annotated to pathways regulating β-cell apoptosis, inflammation, and dedifferentiation. Finally, we identified that the development of hyperglycemia in humans is associated with compromised β-cell BMAL1 expression suggestive of a causative link between circadian clock disruption and β-cell failure in diabetes.

---

The circadian system is a critical component of homeostasis, permitting time-dependent regulation of numerous essential cellular and physiological functions and processes. At the molecular level, circadian rhythms are generated by a molecular clock comprised of transcriptional activators BMAL1 (encoded by the gene *Arntl*) and its heterodimer CLOCK, and repressor genes that encode period (PER1, 2, 3) and cryptochrome (CRY1, 2) proteins^1^. Secondary regulatory loops involving nuclear receptors Rev-Erbα/β and RORα/β provide additional molecular control by acting as respective transcriptional repressors and activators of Bmal1^2,3^. The BMAL1:CLOCK heterodimer is essential for the generation of transcriptional circadian rhythms through DNA binding to conserved motifs along with concurrent recruitment of cell-specific enhancers, transcriptional co-activators, and histone-modifying enzymes essential for the regulation of circadian-regulated target genes^1,4,5^.

Accumulating evidence suggests that a functional circadian system is a critical component of *in vivo* glycemic control^6-8^. Consequently, disruption of normal circadian rhythms results in glucose intolerance and associated β-cell dysfunction^9-12^. Consistently, genetic loss-of–function studies have demonstrated that β-cell circadian clocks are essential for proper regulation of insulin secretion, β-cell postnatal maturation, turnover, and response to diabetogenic stressors^13-15^. More recently, BMAL1:CLOCK-controlled genes in the β-cell have been shown to permit circadian control of insulin secretion through the epigenetic regulation of genes controlling key aspects of intracellular metabolic signaling and insulin exocytosis^16,17^. Despite an increased appreciation for the role of circadian clocks in β-cells, little is known about the contributions of clock disruption to β-cell failure in diabetes.

Progression of both Type 1 (T1DM) and Type 2 (T2DM) diabetes mellitus is associated with alterations in the islet microenvironment, a component of which is the exposure to a subset of key pro-inflammatory cytokines^18,19^. In T1DM diabetes, β-cells are primarily exposed to interleukin (IL-) 1β, tumor necrosis factor α (TNFα) and interferon–γ (IFNγ), as a result of the autoimmune assault and subsequent focal cytokine release mediated by monocytes, macrophages, and T cells^19^. In T2DM, islet cytokine exposure is a result of the complex interplay between 1) adipose-derived circulating cytokines (*e.g.* IL-1β and IL-6)^20,21^, 2) IL-1β release through islet-associated macrophages^22,23^, and 3) islet cell-derived cytokine production mediated through the intracellular stress response^24^. Importantly, acute exposure of islets to pro-inflammatory cytokines recapitulates many aspects of β-cell dysfunction observed during the evolution of diabetes development (*e.g.* loss of first-phase insulin secretion) and leads to alterations in transcriptional networks spanning key pathways regulating β-cell metabolism, insulin biosynthesis, and secretion^25,26^. More importantly, recent studies suggest an important reciprocal relationship between inflammatory responses and the regulation of the core molecular circadian clock^27^. Particularly, evidence of inflammation (NF-κB)-mediated control of core circadian clock genes^28^ and the potential for transcriptional reprogramming^29^ suggests that islet inflammation may promote β-cell failure through its effects on the circadian clock.

In this report, we provide first evidence that pro-inflammatory cytokines disrupt the functionality of the β-cell circadian clock. This process is mediated in part through impaired expression of the key circadian clock transcription factor BMAL1, which was evident at the level of *Bmal1* promoter activation, mRNA, protein, as well as at the level of genome-wide BMAL1 DNA binding patterns. Moreover, we also identified that development of hyperglycemia in humans is associated with loss of β-cell BMAL1 expression suggestive of causative link between inflammation, circadian clock disruption, and β-cell failure in diabetes.

## RESULTS

### *In vitro* exposure to IL-1β disrupts the β-cell circadian clock

Detailed tracking of cell bioluminescence with a clock gene luciferase fusion construct (*e.g. Per*2) permits longitudinal monitoring and assessment of the oscillatory function of peripheral circadian clocks^30^. Thus, to assess the effects of pro-inflammatory cytokines on β-cell circadian clock, we first used *Per2:*LUC^30^ reporter mice crossbred with mice expressing Enhanced Green Fluorescent Protein (GFP) under the control of the insulin promoter^31^ (*Per2*:LUC-MIP:GFP), which allows for examination of circadian activation of the *Per2* promoter in β-cells using real-time bioluminescence tracking (Fig. 1a). Pancreatic islets were isolated from adult (8-12 weeks) *Per2*:LUC-MIP:GFP mice and treated *in vitro* for 72 hours to physiological ranges of key diabetogenic pro-inflammatory cytokines (*e.g.* IL-1β, TNFα, IL-6, and IFNγ)^18,19^. These studies revealed that exposure to IL-1β (0.2-5 ng/ml) led to a significant dampening (∼80%, *p*<.05) of the amplitude and alterations in the phase (peak, 18.6 ± 0.4 vs. 13.8 ± 0.5 hours for IL-1β vs. vehicle, *p*<.05) of *Per2*-driven luciferase oscillations in β-cells (Fig. 1c-e and Supplementary Fig. 1). Importantly, this effect was evident at physiological concentrations previously shown to selectively reduce β-cell function^26^ without a significant decrement in islet cell viability (∼93%, Supplementary Fig. 2). This effect was also IL-1 receptor mediated, as pretreatment with a specific IL-1 receptor antagonist (IL-1ra) rescued alterations in *Per2*-driven bioluminescence in islets (Supplementary Fig. 3). Interestingly, the suppressive effect of IL-1β on the β-cell clock was in contrast to TNFα, IL-6, and IFNγ, all of which showed no significant effect on circadian activation of *Per2*-driven bioluminescence (Fig. 1c-e).

**Figure 1.**
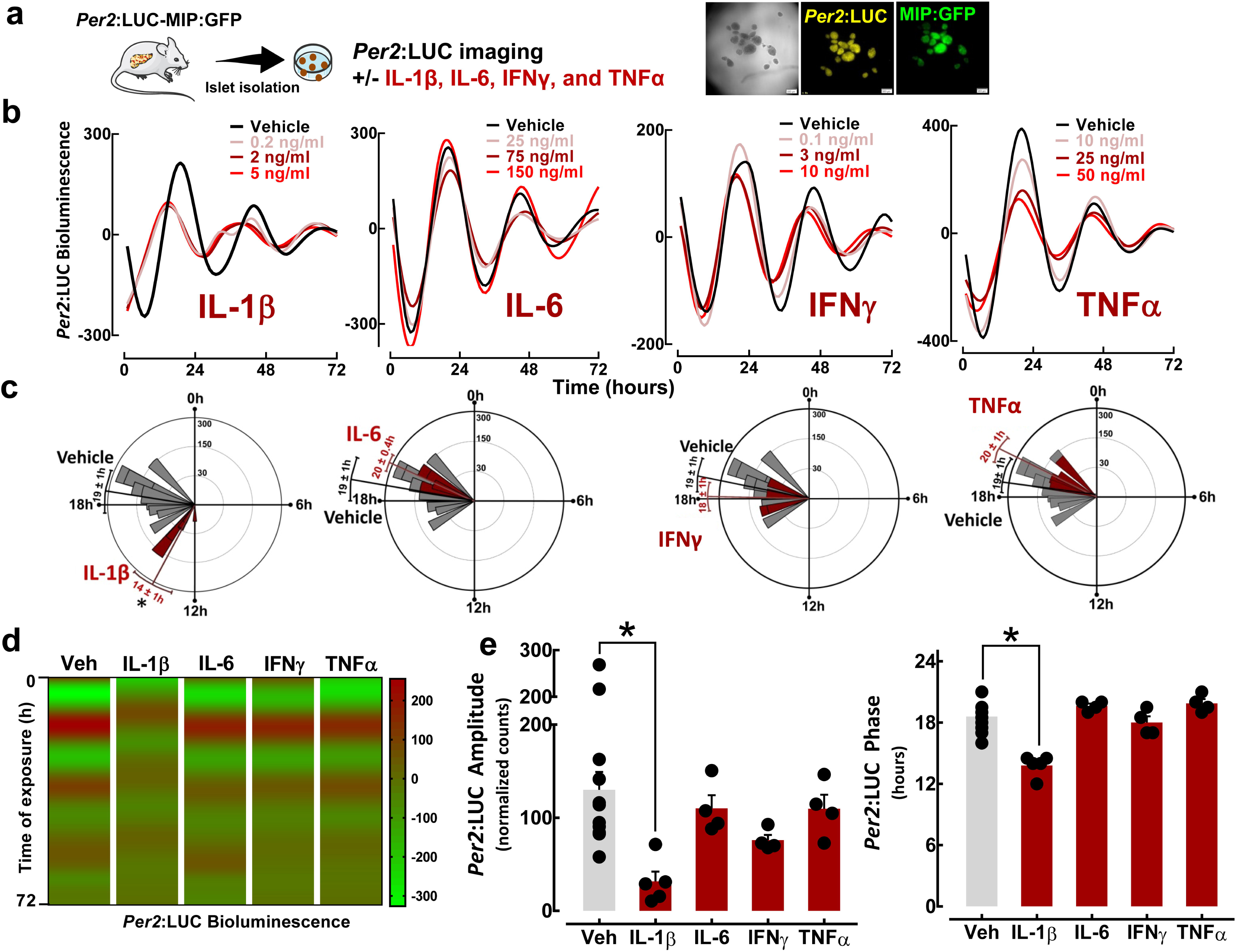
Effects of pro-inflammatory cytokines on β-cell circadian clock elucidated through bioluminescence recordings of islets isolated from *Per2*:LUC-MIP:GFP mice. (a) Diagrammatic representation of the study design indicating that pancreatic islets were isolated from *Per2*:LUC-MIP:GFP mice to assess *Per2*-driven bioluminescence in response to increasing concentrations of IL-1β, TNFα, IL-6, and IFNγ. (b) Representative examples of circadian *Per2*-driven bioluminescence rhythms in batches of 10-15 islets isolated from *Per2*:LUC-MIP:GFP male and female mice exposed for 72 hours to either IL-1β (0.2-5 ng/ml), TNFα (10-50 ng/ml), IL-6 (25-150 ng/ml), IFNγ (0.1-10 ng/ml) vs. vehicle. (c) Rayleigh vector histograms were visualized using Oriana 4.0 (Kovach Computing Services, Anglesey, UK) to assess the significance of the amplitude as well as the phase clustering of peak *Per2*:LUC expression relative to circadian time in response to IL-1β (2 ng/ml), TNFα (25 ng/ml), IL-6 (75 ng/ml), IFNγ (3 ng/ml) or vehicle. The length of each filled vector represents the amplitude of *Per2*-driven bioluminescence, while direction (time) denotes the phase of peak *Per2*-driven bioluminescence in islets cultured with vehicle or the given cytokine condition (n=4-7 independent experiments). The direction of the arrow represents the mean phase vector and the width of the arrowhead represents the variance (standard error) of the mean phase vector. (d) Representative heatmaps displaying bioluminescence traces for islets isolated from *Per2*:LUC-MIP:GFP mice exposed for 72 hours to either IL-1β (0.2-5 ng/ml), TNFα (10-50 ng/ml), IL-6 (25-150 ng/ml), IFNγ (0.1-10 ng/ml) versus vehicle. (e) Bar graphs represent mean amplitude (left) and phase (right) of *Per2*-driven bioluminescence in β-cells exposed to IL-1β (2 ng/ml), TNFα (25 ng/ml), IL-6 (75 ng/ml), IFNγ (3 ng/ml) or vehicle. Values are mean ± SEM (*n*=4-7 independent experiments) and \**p* <.05 denotes statistical significance vs. vehicle.

**Figure 2.**
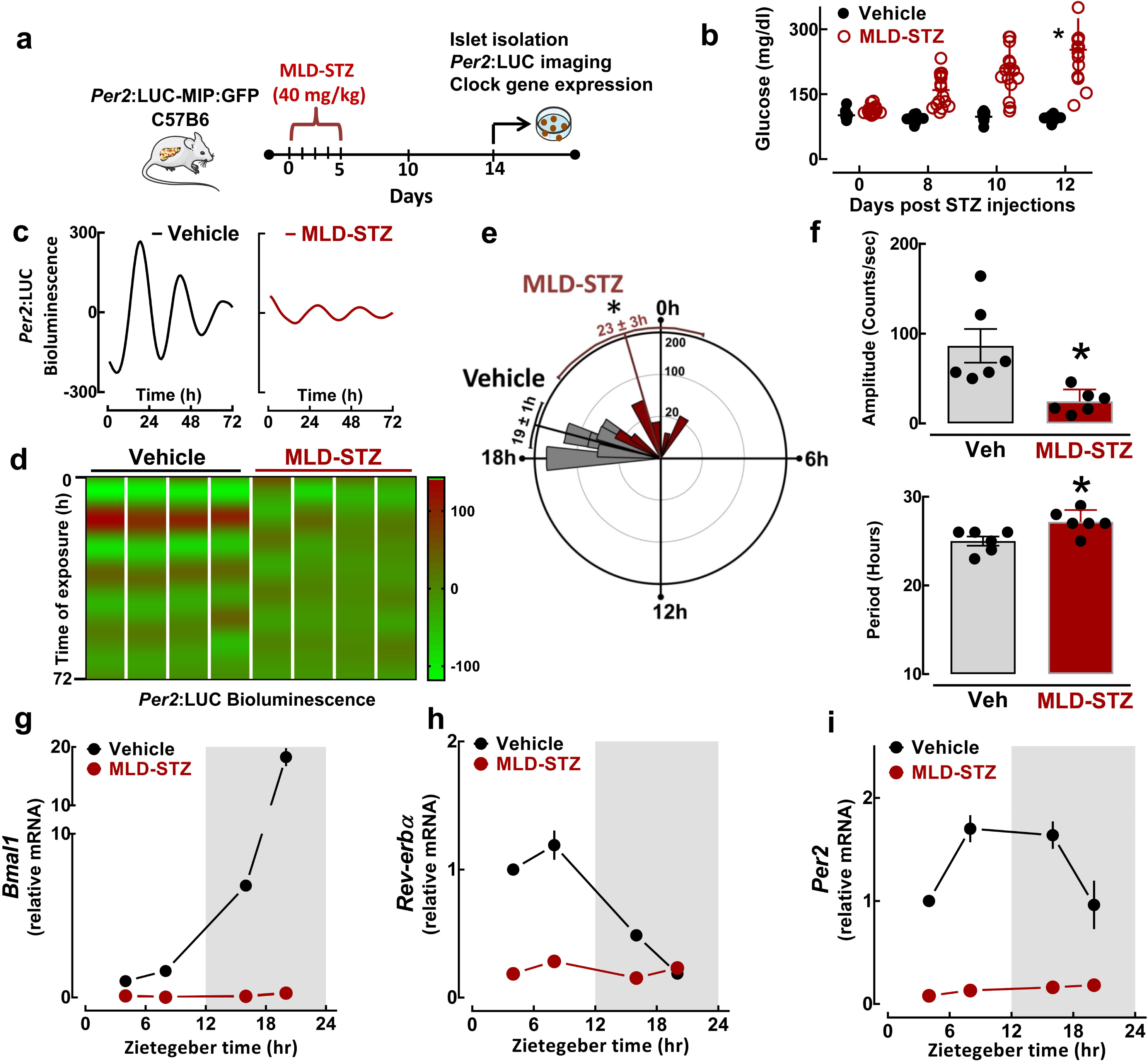
Effects of multiple low dose streptozotocin (MLD-STZ) induced β-cell failure and diabetes on the β-cell circadian clock. (a) Diagrammatic representation of the study design indicating that a subset of *Per2*:LUC-MIP:GFP and C57B6 mice received intraperitoneal injections of streptozotocin (STZ; 40 mg/kg/day) vs. vehicle on five consecutive days to induce pro-inflammatory β-cell failure and diabetes which was confirmed (b) by measurements of blood glucose concentrations post STZ injections. (c) Representative examples of circadian *Per2*-driven bioluminescence rhythms in batches of 10-15 islets isolated from *Per2*:LUC-MIP:GFP mice exposed *in vivo* to MLD-STZ (red) versus vehicle (black). (d) Representative heatmaps displaying bioluminescence traces for islets isolated from *Per2*:LUC-MIP:GFP mice exposed *in vivo* to MLD-STZ (red) versus vehicle (black). Each column represents an independent experiment. (e) Rayleigh vector histograms visualized to assess the significance of the amplitude and the phase clustering of peak *Per2*:LUC expression relative to circadian time in response to MLD-STZ versus vehicle (n=6 independent experiments). (f) Bar graphs represent mean amplitude (top) and circadian period (bottom) of *Per2*-driven bioluminescence in β-cells exposed to MLD-STZ (red) or vehicle (black) (*n*=6 independent experiments). (g-i) Normalized *Bmal1, Rev-Erb*α, and *Per2* mRNA expression obtained from islet lysates of MLD-STZ (red) and vehicle (black) mice collected during ZT 4, 8, 16 and 20 time points in the 24 h circadian cycle. Values are mean ± SEM (*n*=3-4) fold change with vehicle at ZT 4 set as 1. All values are mean ± SEM and \**p*<.05 denotes statistical significance vs. vehicle.

**Figure 3.**
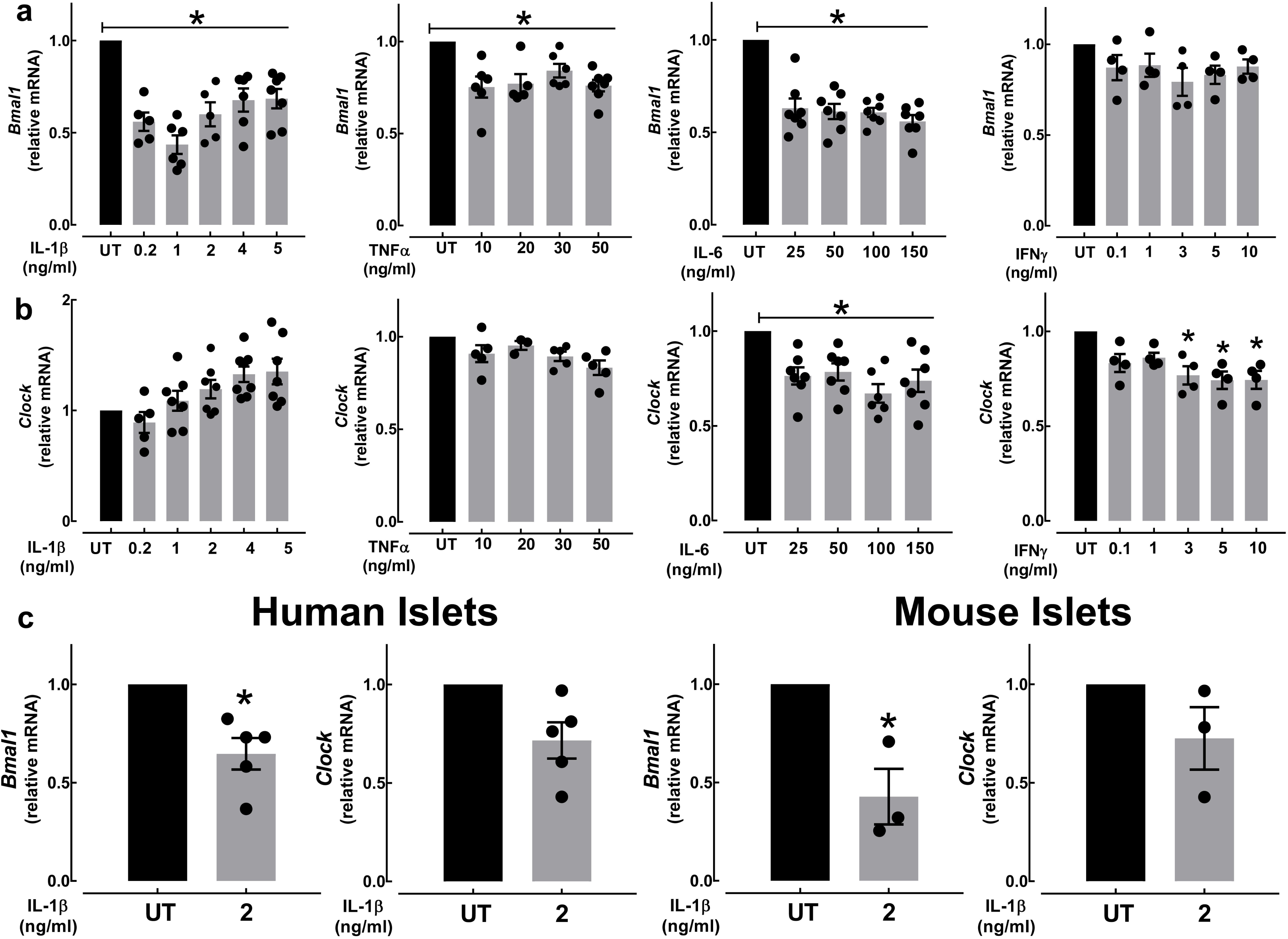
Effects of pro-inflammatory cytokines on *Bmal1* and *Clock* mRNA expression in β-cells and isolated human and mouse islets. (a-b) Normalized *Bmal1* and *Clock* mRNA expression in INS-1 832/13 β-cells exposed for 24 h to either IL-1β (0.2-5 ng/ml), TNFα (10-50 ng/ml), IL-6 (25-150 ng/ml), and IFNγ (0.1-10 ng/ml). Values are mean ± SEM (*n*=4-7 independent experiments per given concentration) and \**p*<.05 denotes statistical significance vs. untreated (UT). (c) Normalized *Bmal1* and *Clock* mRNA expression in isolated human and mouse (C57B6: 8-12 weeks old) islets exposed for 24 h to IL-1β (2 ng/ml) versus untreated (UT). Human islet data represents n=5 independent non-diabetic human islet shipments. Mouse islet data represents n=3 independent experiments. Values are mean ± SEM and \**p*<.05 denotes statistical significance vs. untreated (UT).

### Exposure to inflammation *in vivo* disrupts β-cell circadian clock

To extend the validity of our findings to more physiological conditions, we next investigated β-cell circadian clock function under diabetogenic pro-inflammatory stress *in vivo*. We assessed the β-cell circadian clock in islets isolated from *Per2*:LUC-MIP:GFP mice treated with multiple low dose streptozotocin (MLD-STZ), a mouse model which recapitulates IL-1β–mediated pro-inflammatory β-cell failure in diabetes^32^ (Fig. 2a). To recapitulate aspects of β-cell failure in evolving T2DM in humans, all experiments involving MLD-STZ islets were preformed 14 days post final STZ injection at which point islets displayed preserved islet integrity, but attenuated (∼50%) insulin expression, insulin content, and function (Supplementary Fig. 4, 5). Consistent with the *in vitro* observations, induction of MLD-STZ resulted in disrupted circadian clock function characterized by significant alterations in the amplitude, phase, and the period of *Per2*-driven luciferase oscillations in β-cells (*p*<.05 for all parameters, Fig. 2c-f). Consistently, quantitative RT-PCR performed on islets isolated at multiple time points during the day/night cycle demonstrated impaired expression of key circadian clock genes (*Bmal1, Rev-Erb*α, and *Per2*) in MLD-STZ compared to control mice (*p*<.05 vs. vehicle, Fig. 2g-i). Finally, to confirm whether IL-1β and MLD-STZ disrupts circadian regulation of β-cell function, we assessed self-autonomous circadian rhythms in glucose-stimulated insulin secretion (GSIS) in isolated islets, which was recently shown to be regulated by the β-cell circadian clock^16^. Consistent with previous studies^16^, we observed robust time-dependent rhythms in GSIS following forskolin synchronization of control islets (Supplementary Fig. 5). Importantly, acute treatment with IL-1β (2 ng/ml, 24 h) or *in vivo* exposure to MLD-STZ, abrogated rhythmic regulation of GSIS consistent with the inhibitory effects of IL-1β and MLD-STZ on the β-cell circadian clock (Supplementary Fig. 5).

**Figure 4.**
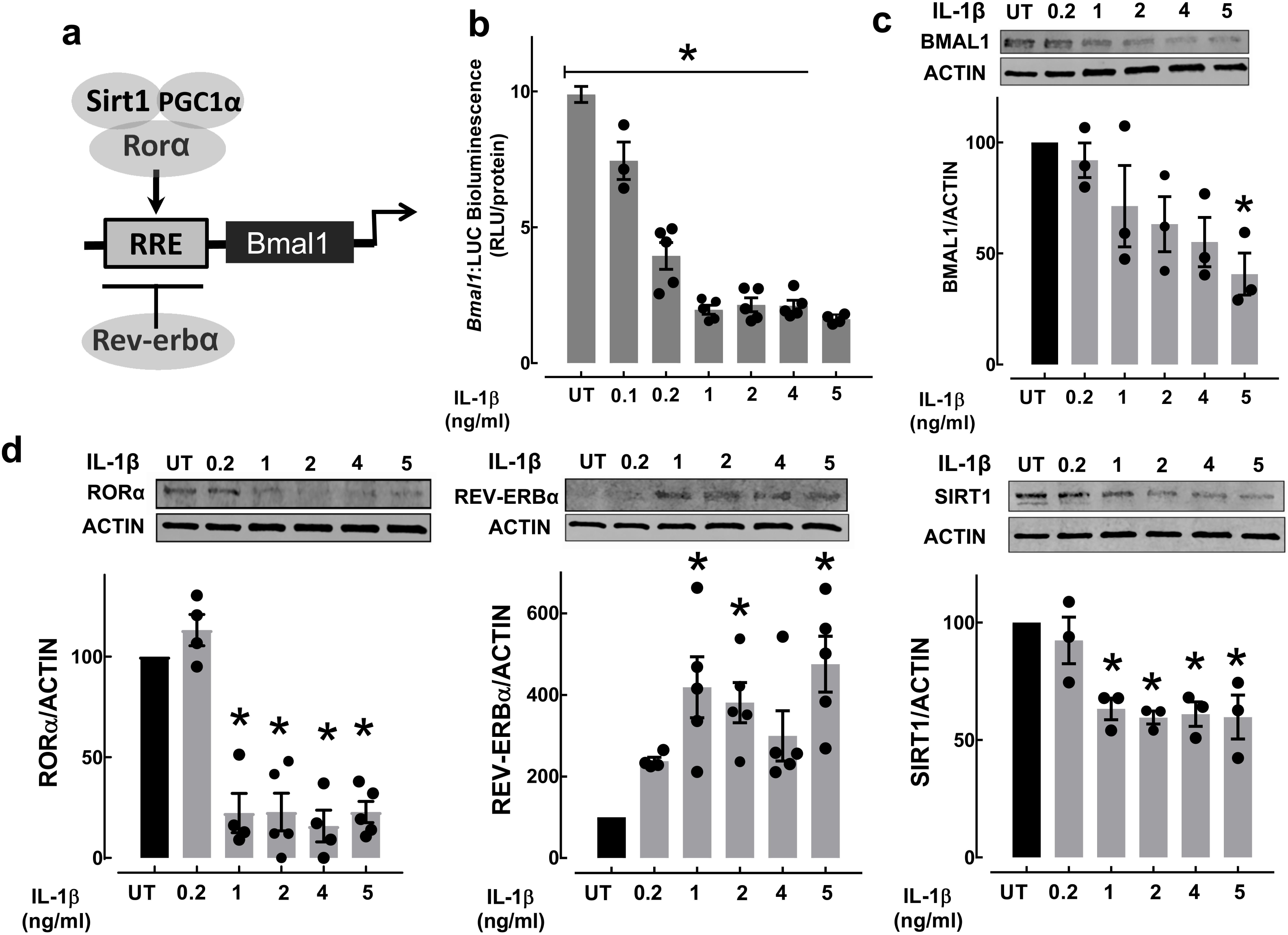
Effects of IL-1β on activators and repressors of *Bmal1* transcription in β-cells. (a) A schematic representation of *Bmal1* promoter regulation. Bmal1 expression is regulated by opposing activities of the orphan nuclear receptors Rorα and Rev-erbα. SIRT1 is a regulator of *Bmal1* transcription mediated in part through cooperative binding with RORα and PGC1α. (b) *Bmal1* promoter activity was assessed using stably-transfected *Bmal1*:LUC reporter expressing INS-1 832/13 β-cells exposed for 24 h to IL-1β (0.1-5 ng/ml). Values are mean ± SEM (*n*=3-6 independent experiments per given concentration) and \**p*<.05 denotes statistical significance vs. untreated (UT). (c-d) BMAL1, RORα, REV-ERBα and SIRT1 protein expression in INS-1 832/13 β-cells exposed for 24 h to IL-1β (0.2-5 ng/ml). Values are mean ± SEM (*n*=3-6 independent experiments per given concentration) and \**p*<.05 denotes statistical significance vs. untreated (UT).

**Figure 5.**
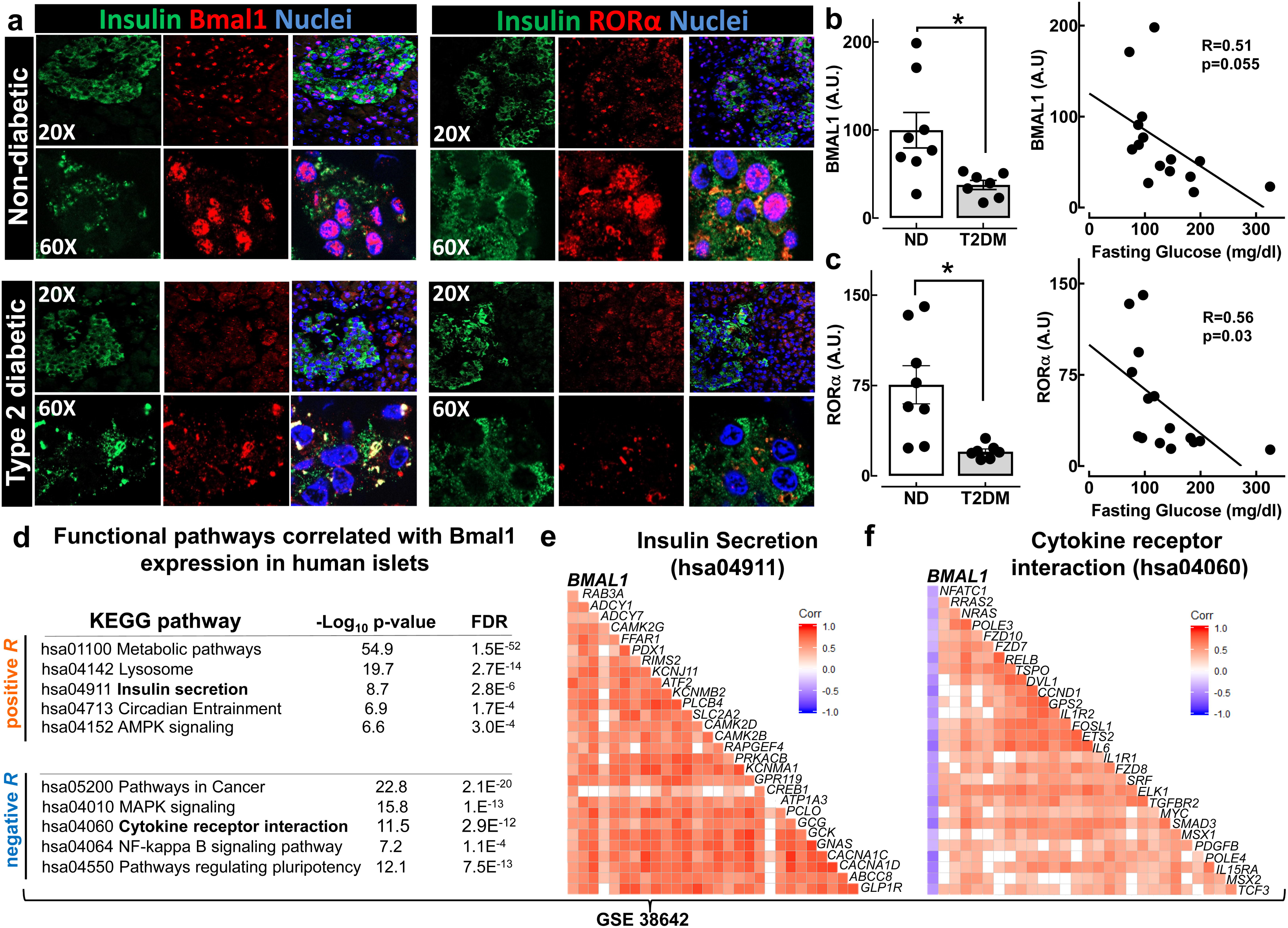
BMAL1 and RORα expression is decreased in human diabetic β-cells. (a) BMAL1 and RORα protein levels assessed by immunofluorescence in human pancreatic tissue obtained at autopsy from nondiabetic subjects and subjects with type 2 diabetes. (b-c) Mean staining intensity for BMAL1 (top) and RORα (bottom) of β-cells from nondiabetic subjects (ND) and subjects with type 2 diabetes mellitus (T2DM). Data are expressed as mean ± SEM (n=7-8 per group) and \**p*<.05 denotes statistical significance vs. non-diabetic (ND). The expression levels of BMAL1 and RORα are negatively correlated with fasting glucose concentrations in individual human subjects (Pearson’s correlation with fasting glucose taken within 1 year of autopsy). (d) Correlation analysis with gene expression data from human pancreatic islets (data set GEO: GSE38642) was performed using Pearson’s correlation analysis. Genes whose expression significantly correlated (r = 0.5 to 1; FDR<0.05) with the expression of *Bmal1* were subjected to KEGG enrichment analysis relevant significantly enriched KEGG pathways shown. (e-f) Pearson’s correlation of *Bmal1* to genes regulating insulin secretion (hsa04911; e) and cytokine receptor interaction (hsa04060; f) is represented by a correlation matrix. Positive and statistically significant Pearson’s correlation coefficients are represented by a red square, while negative coefficients are represented by a blue square. Correlations that were not statistically significant are represented by a white square.

### IL-1β impairs *Bmal1* expression and promoter activation in β-cells

In β-cells, BMAL1:CLOCK heterodimer is required for the generation of circadian rhythms of insulin secretion through transcriptional regulation of genes regulating insulin exocytosis and various aspects of β-cell metabolism and functionality^16^. Indeed, β-cell-specific deletion of *Bmal1* leads to abrogation of circadian rhythms of insulin secretion, glucose intolerance, and recapitulation of numerous aspects of β-cell failure in diabetes^14-16^. Therefore, we next examined the impact of IL-1β exposure (and other pro-inflammatory cytokines) on the mRNA expression of *Bmal1* and *Clock* in the INS-1 832/13 β-cell line in addition to isolated mouse and human islets. Consistent with our previous results, exposure to IL-1β suppressed *Bmal1* mRNA in β-cells (∼60%, *p*<.05, Fig. 3a), but had no suppressive effect on *Clock* expression (Fig. 3b). Interestingly, other diabetogenic cytokines, namely IL-6 and TNFα also showed modest repression of *Bmal1* mRNA in β-cells (*p*<.05, Fig. 3a), whereas exposure to IL-6 and IFNγ led to suppression of *Clock* (*p*<.05, Fig. 3b). Importantly, the suppressive effect of IL-1β on *Bmal1* mRNA was reproduced in primary mouse and human islets (*p*<.05, Fig. 3c) suggesting that the deleterious effects of pro-inflammatory cytokines on the β-cell circadian clock are mediated in part through IL-1β effects on *Bmal1* transcription.

To confirm this we next assessed *Bmal1* promoter activity using a stably-transfected *Bmal1*:luciferase (*Bmal1*:LUC) reporter INS-1 832/13 cell line. Consistent with mRNA expression studies, IL-1β (0.1-5 ng/ml) showed robust dose-dependent suppression of Bmal1 promoter activation (up to 80%, Fig. 4b) which corresponded with decreased BMAL1 protein levels (Fig. 4c). *Bmal1* expression is regulated by opposing activities of the orphan nuclear receptors RORα and Rev-Erbα which respectively play a role as activators and repressors of *Bmal1* transcription through binding to RORE elements on the *Bmal1* promoter^2,3^ (Fig. 4a). Exposure to IL-1β resulted in a substantial decline of RORα levels (∼80%, Fig. 4d) and an induction of REV-ERBα expression (*p*<.05, Fig. 4d). Previous studies have identified an NAD+ dependent deacetylase, Sirt1, as a critical regulator of *Bmal1* transcription mediated in part through cooperative binding with RORα and PPARγ co-activator 1α (PGC1α)^33-35^. In addition, *Sirt1* expression has been suggested to be negatively regulated by inflammation in pancreatic β-cells *in vitro*^36^ and repressed under condition of pro-inflammatory β-cell failure *in vivo*^37^. Consistent with these observations, SIRT1 levels were also significantly repressed upon acute exposure to IL-1β in β-cells (*p*<.05, Fig. 4d), whereas the deleterious effects of IL-1β on Bmal1 promoter activity in β-cells were reversed upon pre-treatment with the Sirt1 activator, resveratrol (Supplementary Fig. 6). Taken together these data suggest that IL-1β suppresses Bmal1 transcription and protein levels by promoting the inhibition of BMAL1 transcriptional activators RORα and SIRT1 and enhancing expression of the transcriptional repressor REV-ERBα.

**Figure 6.**
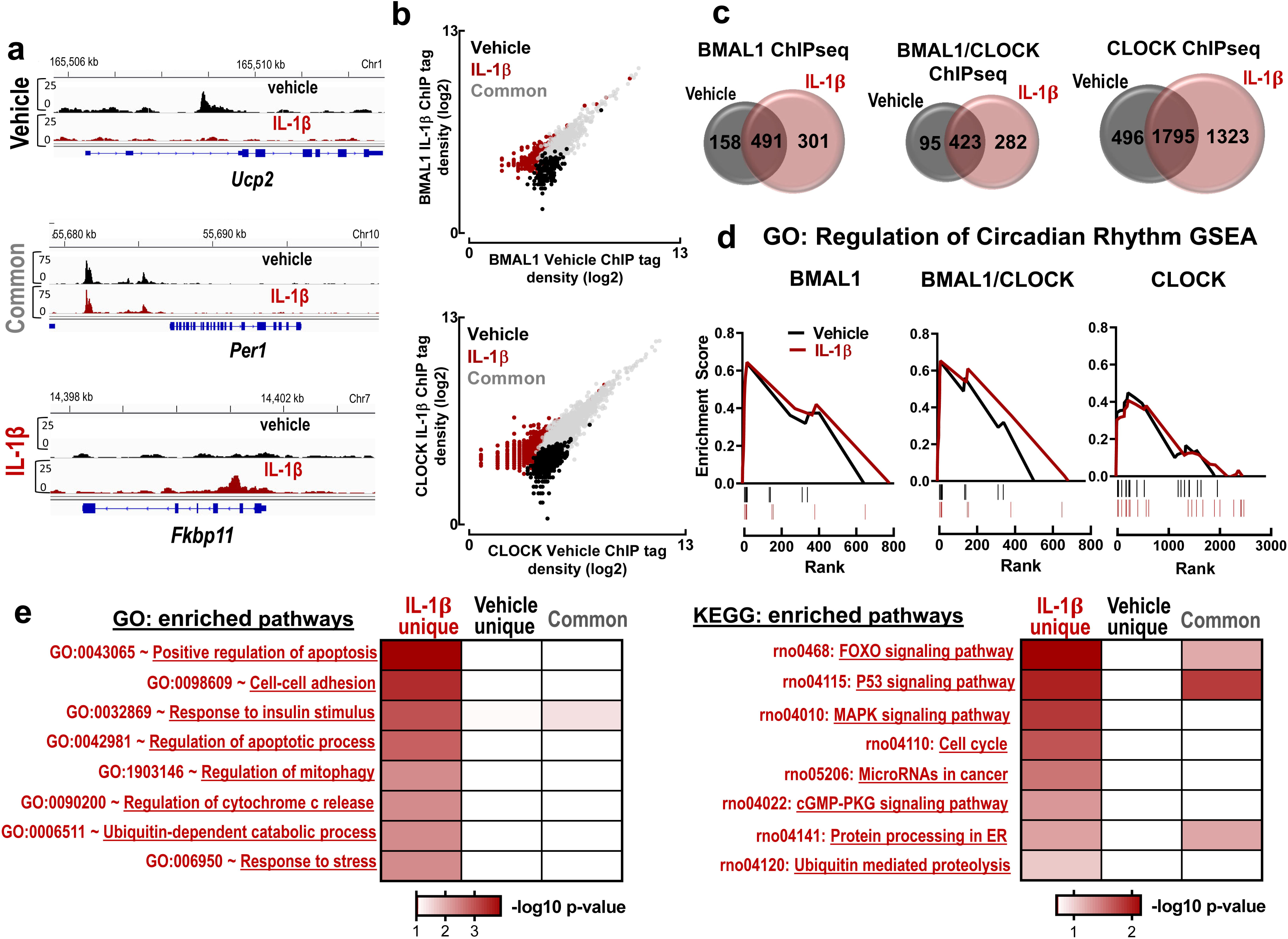
Effects of IL-1β on genome-wide DNA binding patterns of BMAL1 and CLOCK in β-cells. (a) Examples of genome browser view of BMAL1 ChIP-seq binding peaks at *Ucp2* (unique to vehicle conditions), *Per1* (common for vehicle and IL-1β conditions) and *Fkbp11* (unique to IL-1β conditions) gene loci. (b) Scatter plots represent ChIP-seq tag density distribution for BMAL1 and CLOCK in INS-1 832/13 β-cells exposed to either IL-1β (2 ng/ml) or vehicle control. Grey dots indicate binding peaks shared between vehicle and IL-1β conditions, whereas black and red dots indicate tag density distribution unique to vehicle or IL-1β respectively. (c) Venn diagrams indicate an overlap and unique binding peaks for BMAL1, CLOCK and BMAL1:CLOCK cobound for vehicle and/or IL-1β conditions. (d) Gene-set enrichment analysis (GSEA) performed using pre-ranked analysis of target peak size of annotated binding sites in vehicle and IL1-β conditions (unique plus common) using GO biological process “Regulation of Circadian Rhythm” gene set [GO term: 0042752]. FDR q = 0.00. (f) Functional gene ontology (GO) and pathway (KEGG) enrichment analysis of the genomic sites co-occupied by BMAL1:CLOCK exclusive to the IL-1β treatment conditions, *p*<.05.

### Diabetes in humans is associated with impaired β-cell expression of key circadian transcriptional regulators

To address whether the development of diabetes in humans is associated with impaired β-cell expression of key circadian transcriptional regulators, we first performed detailed immunofluorescence analysis of Mayo Clinic autopsy-derived human pancreatic tissue collected from subjects with T2DM vs. age/BMI-matched non-diabetic controls (Fig. 5, Supplementary Table 2). Consistent with our previous observations, nuclear expression of both BMAL1 and RORα was decreased specifically in β-cells of patients with T2DM (*p*<.05 vs. non-diabetics, Fig. 5a-c and Supplementary Fig. 7). Furthermore, β-cell BMAL1 (and RORα) expression demonstrated a robust negative correlation with T2DM patient fasting plasma glucose levels indicative of a plausible relationship between β-cell circadian clock and glycemic control in humans (Fig. 5a-c). In addition, we also performed islet transcriptome bioinformatics reanalysis of an independent dataset (GEO: GSE38642) containing human islet genome-wide mRNA expression from non-diabetic, pre-diabetic, and T2DM individuals^38,39^. Interestingly, *Bmal1* mRNA expression in human islets across the non-diabetic to prediabetic/T2DM range demonstrated a positive correlation with the expression of transcripts functionally annotated to KEGG pathways controlling circadian rhythms and β-cell function (*e.g.* circadian entrainment, insulin secretion, metabolic pathways, etc., Fig. 5d-f). In contrast, *Bmal1* mRNA expression in human islets was negatively correlated with KEGG pathways involved in the regulation of β-cell failure, inflammation, and dedifferentiation (*e.g.* MAPK signaling, NF-κB signaling, pathways in cancer etc., Fig. 5d-f). This data provides first evidence of impaired BMAL1 (and RORα) expression in human β-cells in diabetes which correlates with the expression of signaling pathways involved in the regulation of β-cell function and failure.

### IL-1β augments DNA binding patterns of BMAL1 and CLOCK in β-cells

Recent reports demonstrate that the effects of cellular inflammatory stress may extend to reprogramming of BMAL1:CLOCK genomic binding toward a distinct circadian transcriptional program^29,40,41^. Therefore, we next set out to determine whether exposure to IL-1β also alters genome-wide DNA binding patterns of BMAL1 and CLOCK in β-cells. Chromatin immunoprecipitation followed by sequencing (ChIP-seq) of INS-1 832/13 β-cells revealed a subset of BMAL1 and CLOCK binding sites which were unique (or common) for vehicle or IL-1β-stimulated conditions (Fig. 6a-b). Interestingly, IL-1β treatment resulted in an overall increase in total binding sites for BMAL1 (∼18% increase), CLOCK (27%) and BMAL1:CLOCK cobound (27%), indicative of a wider distribution of BMAL1 and CLOCK binding in β-cells during IL-1β-mediated inflammation (Fig. 6c and Supplementary Fig. 8). First, to test whether IL-1β leads to relocalization of BMAL1/CLOCK binding from sites annotated to genes regulating circadian rhythms, we performed gene-set enrichment analysis (GSEA) of annotated target genes in vehicle and IL1-β conditions (unique plus common) using GO biological process “Regulation of Circadian Rhythm” gene set [GO term: 0042752]. GSEA analysis revealed a significant enrichment for circadian-regulated genes under both vehicle and IL-1β conditions (Fig. 6d, [FDR] q = 0.00), implying that IL-1β doesn’t alter BMAL1/CLOCK genome-wide binding to core circadian clock genes in β-cells. Secondly, we performed functional pathway analysis (GO and KEGG) on a unique subset of 301 (BMAL1), 1323 (CLOCK), and 282 (BMAL1:CLOCK cobound) binding sites exclusive to the IL-1β treatment conditions (Fig. 6e). Importantly, this analysis revealed that exposure to IL-1β results in the enrichment of BMAL1 and CLOCK binding sites in the proximity of genes functionally annotated to pathways regulating β-cell apoptosis, inflammation, and dedifferentiation (*e.g.* mitophagy, FOXO signaling, MAPK signaling etc.) (Fig. 6e). This data suggests that under pro-inflammatory conditions brought on by exposure to IL1-β, the core components of the β-cell circadian clock is, in part, reorganized towards transcriptional regulation of pro-inflammatory and pro-apoptotic pathways, consistent with recent observations of circadian reprogramming in the lung^29^, macrophages^42^ and hepatocytes^23^.

## DISCUSSION

Development of hyperglycemia in both T1DM and T2DM is preceded by β-cell secretory dysfunction progressing toward induction of β-cell apoptosis and/or dedifferentiation^43-45^. Although there are many notable differences in the pathophysiology of β-cell failure between the two forms of diabetes, exposure to pro-inflammatory cytokines plays a prominent role in the pathophysiology of both diseases^46^. Indeed, acute *in vitro* exposure of isolated islets to key pro-inflammatory cytokines (*e.g.* IL-1β, TNFα, IL-6, and IFNγ) recapitulates many aspects of β-cell dysfunction observed during the evolution of diabetes development such as loss glucose-stimulated insulin secretion, induction of oxidative and ER stress response, increased autophagy, and loss of transcriptional identity^26^. Consistently, cytokine exposure of primary β-cells leads to alterations in diverse transcriptional networks spanning hundreds of genes belonging to key pathways regulating β-cell metabolism, insulin biosynthesis and secretion^25^. However, despite an increased understanding into the biological effects of β-cell cytokine exposure, the molecular basis underlying the deleterious effects of pro-inflammatory cytokines on β-cell transcriptional networks are not fully understood.

The current study examined effects of key diabetogenic pro-inflammatory cytokines on the β-cell circadian clock. In particular, among the tested cytokines IL-1β was demonstrated to have the most profound disruptive effect on the expression of core clock transcription factors (*e.g.* Bmal1 and Clock) and clock-mediated circadian transcriptional and physiological oscillations. Indeed, ample evidence describes the involvement of IL-1β in Type 1 diabetes and more recently, Type 2 diabetes^18,19^. In Type 1 diabetes, IL-1β is produced primarily by monocytes and macrophages and exerts a direct effect on β-cell function and survival, often in concert with other pro-inflammatory cytokines^19^. In Type 2 diabetes, the origin of IL-1β exposure to the β-cells is more complex and controversial; however is likely to include a combination of systemic adipose-derived and “local” islet inflammasome/macrophage–derived IL-1β secretion^20-23^. Most notably, IL-1β exposure at submaximal concentrations (and duration) used in our study has been shown to preferentially induce defects in β-cell secretory function by altering key mediators of insulin exocytosis and granular machinery (*e.g.* Snap25), an effect shown to be unique to IL-1β among other pro-inflammatory cytokines^26^. Interestingly, recent studies have shown that self-autonomous circadian rhythms in glucose-stimulated insulin secretion are driven by BMAL1:CLOCK mediated regulation of transcriptional enhancers encoding expression of genes involved in the assembly and translocation of insulin secretory granules^16^. It is thus plausible to hypothesize that the deleterious effects of IL-1β on the β-cell are mediated in part through repression of circadian clock machinery and circadian control of insulin secretion.

Our study shows that IL-1β disrupts β-cell circadian clock in part through interfering with the expression of key circadian transcription factor Bmal1. Indeed, Bmal1 is the only non-redundant clock gene indispensable for the regulation of circadian rhythms^47^. In β-cells, Bmal1 is required for the regulation of functional maturation^48^, insulin secretion^16^, β-cell turnover^15^ and mitochondrial function^14^. Thus, attenuated Bmal1 expression recapitulates many features of β-cell failure in diabetes^49^. IL-1β appears to regulate β-cell Bmal1 expression by promoting inhibition of Bmal1 transcriptional activators RORα and SIRT1 and enhancing expression of the transcriptional repressor REV-ERBα. This is consistent with work that has identified an NAD+ dependent deacetylase Sirt1 as a key regulator of Bmal1 transcription mediated in part through cooperative binding with RORα and PPARγ co-activator 1α (PGC1α)^33-35^. Notably, Sirt1 expression in β-cells is attenuated in response to IL-1β in NF-kB-dependent manner and Sirt1 expression is also dysregulated in β-cells of animal models of T1DM and T2DM^36,37^. Finally, our work also suggests that IL-1β may control β-cell Bmal1 expression by regulating expression of its transcriptional repressor, REV-ERBα. Although the specific mechanisms underlying induction of REV-ERBα in IL-1β-treated β-cells was not delineated, recent reports show that dysregulated expression of a known downstream target of IL-1β (c-Myc) disrupts the circadian clock by inducing REV-ERBα to repress expression of BMAL1^50^.

Current data also suggests that the effects of inflammation on the β-cell circadian clock may extend beyond suppression of core circadian clock genes, but also encompass epigenetic reprogramming toward a distinct circadian transcriptional program as recently demonstrated in lung and liver cells exposed to pro-inflammatory conditions such as endotoxemia, LPS, and high fat diet^29,40,41^. For example, lung endotoxemia led to an emergence of circadian rhythms in genes enriched for key pro-inflammatory signaling pathways^29^. Moreover, in hepatocytes, activation of pro-inflammatory NF-κB signaling resulted in relocalization of BMAL1 genomic sites during LPS-induced inflammation towards genes involved in immune, apoptotic, and metabolic programs^40^. This effect was shown to be dependent on p65 expression and enhanced chromatin accessibility, providing a plausible explanation for the emergence of BMAL1:CLOCK novel binding sites in response to inflammation^40^. Additionally, BMAL1:CLOCK has been recently shown to function as a pioneer transcription factor thereby regulating DNA accessibility of numerous tissue-specific and ubiquitous transcription factors through nucleosome removal and chromatin modification^4^. This model therefore suggests that the activation of BMAL1:CLOCK target genes requires complex chromatin remodeling and recruitment of various transcription factors which may be influenced by changes in cellular environment due to pro-inflammatory stress^5^. Indeed, in β-cells our study shows that exposure to IL-1β leads to the addition of BMAL1 (and CLOCK) genomic binding sites in the proximity of genes annotated to pathways regulating β-cell failure such as apoptosis, inflammation, and dedifferentiation.

Little is known regarding the regulation of circadian clocks in human β-cells in health and during progression to both forms of diabetes. Classic studies by Boden and colleagues have demonstrated *in vivo* circadian oscillations in insulin secretion and glucose tolerance in humans^51^. These early observations are supported by the presence of a robust circadian transcriptional program in isolated human islets that is dependent on core circadian clock gene expression^16,52^. Stamenkovic and colleagues previously reported that isolated human islets from T2DM patients exhibit a significant decrease in the expression of core circadian clock genes *PER1, PER2*, and *CRY2*, but failed to observe reduction in *BMAL1* expression compared to non-diabetic controls^53^. One potential reason for the discrepancy between the findings by Stamenkovic et al. and our observations of reduced BMAL1 protein in T2DM may stem from different T2DM tissues sources in the two studies (e.g. cultured isolated islets vs. autopsy-derived pancreas). Islet inflammation and cytokine exposure in T2DM is a result of the complex interplay between exposure to systemic circulating cytokines^20,21^ and induction of islet pro-inflammatory microenvironment^22,23^. Thus it is plausible that typical islet isolation process and consequent prolonged culture of human T2DM islets may not fully recapitulate islet pro-inflammatory microenvironment commonly observed *in vivo*.

Our data provides the evidence that the development of hyperglycemia in humans is associated with compromised expression of BMAL1 and RORα in human β-cells suggestive of a potential link between circadian clock disruption and β-cell failure in diabetes. Indeed, studies in humans have reported loss of circadian regulation of insulin secretion in diabetes-prone populations^54,55^. Importantly, these effects are likely due to increased islet exposure to pro-inflammatory cytokines, rather than exposure to hyperglycemia *per se*, since culture of islets in hyperglycemic conditions alone fails to alter circadian clock function^9^. Moreover, potential IL-1β-mediated addition of novel BMAL1 (and CLOCK) genomic binding sites in β-cells suggests that inflammation-mediated circadian reprogramming may play a contributory role during evolution of β-cell failure in humans. Future studies will be required to address the role of the β-cell circadian clock in humans during the transition to T1DM and T2DM. In this regard, therapeutic modulation of the circadian system may provide a promising novel approach to treat metabolic disorders such as diabetes mellitus.

## MATERIALS AND METHODS

### Animal models

A total of 34 C57B6 and 35 *Per2*:LUC-MIP:GFP (on C57B6 background) mice at age 8-12 weeks (equal proportions of male/female) were used in the current study. *Per2*:LUC-MIP:GFP mice were generated through breeding of *Per2*:LUC^30^ reporter mice with mice expressing Enhanced Green Fluorescent Protein (GFP) under the control of the insulin promoter^31^. All animals were housed under standard 12 h light, 12 h dark (LD) cycle and provided with ad libitum chow diet (14% fat, 32% protein, and 54% carbohydrates; Harlan Laboratories, Indianapolis, IN). By convention, the time of lights on (0600 h) is denoted as Zeitgeber Time (ZT) 0 and time of lights off (1800 h) as ZT12. A subset of mice received intraperitoneal injections of streptozotocin (STZ; 40 mg/kg/day in citrate buffer, pH 4.5) or vehicle (citrate buffer) on five consecutive days to induce pro-inflammatory β-cell failure and diabetes^32^. All experimental procedures involving animals described were approved by Mayo Clinic Institutional Animal Care Committee.

### Cell models

INS-1 832/13 cells were generously provided by Dr. Christopher Newgard (Duke University, Durham, NC). INS-1 832/13 cells were cultured in RPMI media with L-glutamine, 10% fetal bovine serum (FBS), 1 mM sodium pyruvate, 10 mM HEPES, and 50 µM β-mercaptoethanol. HEK 293T cells were cultured in Dulbecco’s Modified Eagle Medium (DMEM) supplemented with 10% FBS, 50 U/ml penicillin, and 50 µg/ml streptomycin. All cytokines, IL-1β (201-LB-005/CF), IL-6 (206-IL-010/CF), TNFα (210-TA-020/CF), and IFNγ (585-IF-100/CF) were from R&D Systems (Minneapolis, MN). *Bmal1* reporter INS-1 832/13 cell line (*Bmal1*:LUC) was generated by transducing INS-1 832/13 cells with lentiviral vectors expressing firefly luciferase under the *Bmal1* promoter. Transduced cells were purified by puromycin selection. Lentiviral production has been previously described^56^. In brief, vector plasmid pABpuro-BluF (a gift from Steven Brown, Addgene plasmid #46824), and packaging plasmids PEx-QV and pMD-G were co-transfected into HEK 293T cells using Fugene 6 (Roche). After 4 days, supernatants were filtered, and subjected to ultracentrifugation to concentrate the lentiviral vectors.

### Mouse islet isolation, synchronization and measurements of glucose-stimulated insulin secretion

Pancreatic mouse islets were isolated using collagenase method as previously described^57^ and allowed to recover overnight incubated in standard RPMI 1640 medium (11 mM glucose) supplemented with 10% fetal bovine serum at 37°C. To measure circadian rhythms in glucose-stimulated insulin secretion isolated islets were synchronized through 1 hour exposure to 10 μM forskolin^16^. Glucose-stimulated insulin secretion was assessed by static incubation at basal (4 mM glucose per 30 minutes) and hyperglycemic conditions (16 mM glucose per 30 minutes) with insulin measured by ELISA (ALPCO).

### Per2:LUC islet bioluminescence studies

After euthanasia, pancreatic islets were isolated from *Per2*:LUC-MIP:GFP mice using standard collagenase method^57^. *Per2*-driven bioluminescence emanating from individual GFP^+^ islet cells were imaged by cooled intensified charge-coupled device (ICCD) camera outfitted with a temperature and gas controlled incubation chamber (LV-200, Olympus, MN, USA). Batches of 25 islets were placed in individual wells of a Nunc™ Lab-Tek™ Chambered coverglass (155383; ThermoFisher Scientific, Waltham, MA), each containing standard RPMI 1640 medium with 10% fetal bovine serum and 0.1 mM D-luciferin (L-8220, Biosynth, Itaska, IL). The islets were placed in the Luminoview LV200 Bioluminescence Imaging System and bioluminescence was measured for 1 min at intervals of 10 min and continuously recorded for at least 3 days. Bioluminescence signal was quantified from the acquired images using cellSens software (Olympus, Center Valley, PA). Raw data were normalized by subtraction of the 24 h running average from raw data and then smoothed with a 2 h running average. Analysis of circadian rhythm parameters (*i.e.* circadian period, phase angle, and relative and absolute amplitude) was performed using JTK_CYCLE^58^.

### Bmal1 Luciferase Assay

*Bmal1*:LUC cells were seeded in 12 well plates (1 × 10^6^ / well). Cells were incubated with serial dilutions of the cytokines IL-1β, TNFα, IL-6, and IFNγ for 24 hours. Cells were washed in 1X PBS and 1X cell lysis buffer was added. Plates were put on an orbital shaker for 20 minutes, and cells lysates were collected, centrifuged, and the resulting supernatants were assayed. Cell lysates were added to 96 well opaque plates (Costar) in duplicate and ran on a Promega GloMax-Multi+ Detection System. Values were normalized to total protein content measured by bicinchoninic acid (BCA) assay.

### Quantitative real-time PCR analysis

INS-1 832/13 cells were plated in either 6- or 12-well plates at 1-1.5 × 10^6^ cells/well. Cells were treated with different cytokines at the concentrations noted. After treatment for 24 hours, total RNA was isolated using the RNeasy Mini kit (Qiagen). Complement DNA (cDNA) was transcribed from 300-400 ng of RNA with the iScript cDNA Synthesis kit (BioRad) and the resulting cDNA was mixed with SYBR Green Master Mix (ABI) with gene-specific primers (Supplementary Table 1) for a total reaction volume of 25 µl. qRT-PCR analysis was performed using the ABI StepOnePlus Real-Time PCR System and the results were normalized to β-actin expression.

### Western blot analysis

INS-1 832/13 cells were seeded in 6 well plates (1.5 × 10^6^ /well). Upon treatment with cytokines for 24 hours, total protein lysates was isolated using NP-40 buffer (Amresco, J619) supplemented with protease/phosphatase inhibitor cocktail (Cell Signaling Technology, #5872). Protein lysates were ran on Mini-Protean TGX 4-20% gradient SDS-PAGE gels (BioRad) and transferred to 0.2 µm PVDF Trans-blot Turbo membranes (BioRad) using the Trans-blot Turbo System (BioRad). Membranes were blocked in 5% milk (BioRad) and incubated with antibodies for BMAL-1 (Abcam, ab3350), CLOCK (Millipore, AB2203), RORα (Abcam, ab60134), Rev-Erbα (Cell Signaling Technology, #13418), Sirt1 (Abcam, ab12193) or β-actin (Millipore, MAB1501). The corresponding IRDyes (LiCor) were used for the secondary antibodies and images were obtained with a LiCor Odyssey Fc.

### Chromatin Immunoprecipitation sequencing

INS-1 832/13 cells were cross-linked with 1% formaldehyde for 10 min, followed by quenching with 125 mM glycine for 5 min at room temperature. Fixed cells were washed with TBS and the cell pellets were frozen at −80°C. ChIP-seq was performed as previously described^59^. Briefly, cells were resuspended in cell lysis buffer (10 mM Tris-HCl, pH 7.5, 10 mM NaCl, 0.5% NP-40) and incubated on ice for 10 min. The lysates were washed with MNase digestion buffer (20 mM Tris-HCl, pH 7.5, 15 mM NaCl, 60 mM KCl, 1 mM CaCl_2_) and incubated for 20 minutes at 37 °C in the presence of MNase. After adding the same volume of sonication buffer (100 mM Tris-HCl, pH 8.1, 20 mM EDTA, 200 mM NaCl, 2% Triton X-100, 0.2% Sodium deoxycholate), the lysate was sonicated for 5 min (30 sec-on / 30 sec-off) in Diagenode bioruptor and centrifuged at 15,000 rpm for 10 min. The cleared supernatant equivalent to about 40 × 10^6^ cells was incubated with 5 μg of anti-BMAL1 (Abcam, ab3350) or anti-CLOCK (Abcam, ab3517) antibodies overnight. After adding 30 μl of prewashed protein G-agarose beads, the reactions were further incubated for 3 hours. The beads were extensively washed with ChIP buffer, high salt buffer, LiCl_2_ buffer, and TE buffer. Bound chromatins were eluted and reverse-crosslinked at 65°C overnight. DNAs were purified using Min-Elute PCR purification kit (Qiagen) after treatment of RNase A and proteinase K. The enrichment was analyzed by targeted real-time PCR in *Per2* and negative genomic loci. For the next-generation sequencing, ChIP-seq libraries were prepared from 5-10 ng of ChIP and input DNAs with the ThruPLEX^®^ DNA-seq Kit V2 (Rubicon Genomics, Ann Arbor, MI). The ChIP-seq libraries were sequenced to 51 base pairs from both ends using the Illumina HiSeq 4000 in the Mayo Clinic Center for Individualized Medicine Medical Genomics Facility. Raw sequencing reads were processed and analyzed using the HiChIP pipeline^60^ to obtain visualization files and a list of peaks. Briefly, paired-end reads were mapped to the rat reference genome (release rn6) by BWA^61^ with default settings, and only uniquely mapped reads remained for further analysis. Peaks were called using the MACS2 algorithm^62^ for CLOCK and BMAL1 both at FDR <=1%. Venn-diagram and correlation analysis were performed by our in-house python scripts. Functional enrichment analysis was performed using DAVID to determine enrichment of Biological Process terms and KEGG (Kyoto Encyclopedia of Genes and Genomes) pathways of identified gene targets.

### Human pancreas and islet studies

Human pancreas was procured from the Mayo Clinic autopsy archives with approval from the Institutional Research Biosafety Board. Paraffin-embedded pancreatic sections were immunostained for Insulin (ab7842; Abcam), Bmal1 (ab93806; Abcam), Rorα (ab60134; Abcam) with Vectashield-DAPI mounting medium (Vector Laboratories). Blinded slides were viewed, imaged, and analyzed using a Zeiss Axio Observer Z1 microscope (Carl Zeiss Microscopy, LLC, NY, USA) and ZenPro software (Carl Zeiss Microscopy, LLC.). ImageJ (National Institutes of Health, Bethesda, MD) was used to quantify BMAL1 and RORα expression by measuring average pixel intensity within insulin-positive β-cells, following background subtraction^63^. Isolated human islets were obtained from Prodo Laboratories (Irvine, CA).

### Statistical analysis and calculations

Statistical analysis was performed using ANOVA with post hoc tests wherever appropriate (GraphPad Prism v.6.0, San Diego, CA, USA). Rayleigh vector histograms representing the amplitude and timing of the peak *Per2*:LUC bioluminescence was visualized using Oriana 4.0 (Kovach Computing Services, Anglesey, UK). All data was presented as means ± SEM and assumed statistically significant at *p*<.05.

## ACKNOWLEDGMENTS

We acknowledge funding support from the National Institutes of Health (R01DK098468 to AVM and T32-HL105355 to NJ) and the Center for Regenerative Medicine (Mayo Clinic, Rochester, MN).

## AUTHOR CONTRIBUTIONS

N.J. contributed to study design, conducted experiments, assisted with the data analysis, interpretation and preparation of the manuscript. M.B. conducted experiments, assisted with the data analysis, interpretation and preparation of the manuscript. K.R. contributed to study design, conducted experiments, and reviewed the manuscript. T.H. conducted experiments, and reviewed the manuscript. Z.Y. contributed to data analysis and interpretation, and reviewed the manuscript. J.H. conducted experiments, assisted with the data analysis and reviewed the manuscript. T.O. contributed to study design, data analysis and interpretation and preparation of the manuscript. A.V.M. designed, interpreted the studies, and wrote the manuscript.

A.V.M is the guarantor of this work and, as such, had full access to all data in the study and takes responsibility for the integrity and accuracy of data analysis.

