## Supplementary Table 1 for "Pro-inflammatory cytokines disrupt β-cell circadian clocks in diabetes"

| Species | Forward (5’ to 3’) | Reverse (5’ to 3’) |
| --- | --- | --- |
| Rat |  |  |
| *Bmal1* | TCCGATGACGAACTGAAACA | CGGTCACATCCTACGACAAA |
| *Clock* | CAGCCGCATCCTTCAGTT | CATGGAGCAACCGAGATGT |
| *Reverbα* | CAGCGAGAAGCTCAACTCTCTG | CCATTCCCGAGCGGTCTGC |
| *Rorα* | TGCCACCTACTCCTGTCCTC | CCGTCCATGTAGGTGCTGAG |
| *Actin* | ACTCCTACGTGGGCGACGAGG | CAGGTCCAGACGCAGGATGGC |
| Human |  |  |
| *Bmal1* | ACGGTGGTGCTGGCTAGAGTGTA | CTTCCAAAAACCACAGTGCCCGGC |
| *Clock* | CCAGCCACCGCAACAATT | GGATTCCCATGGAGCAACCTA |
| *Gapdh* | GAAGGTGAAGGTCGGAGTC | GAAGATGGTGATGGGATTTC |
| Mouse |  |  |
| *Bmal1* | TCAAGACGACATAGGACACCT | GGACATTGGCTAAAACAACAGTG |
| *Clock* | CAAAATGTCACGAGCACTTAATGC | ATATCCACTGCTGGCCTTTGG |
| *Per2* | ATGCTCGCCATCCACAAGA | GCGGAATCGAATGGGAGAAT |
| *Reverbα* | TGAACGCAGGAGGTGTGATTG | GAGGACTGGAAGCTATTCTCAGA |
| *Gapdh* | AGGTCGGTGTGAACGGATTTG | TGTAGACCATGTAGTTGAGGTCA |
