## Supplementary Table 2 for "Pro-inflammatory cytokines disrupt β-cell circadian clocks in diabetes"

Clinical characteristics of autopsy pancreas study subjects

| | Age (years) | BMI | Fasting Glucose (mg) |
|---|---|---|---|
| Non-diabetic | 74±4 | 25±1 | 93±5 |
| Type 2 diabetic | 75±5 | 36±2* | 188±25* |

*$P<0.05$ denotes statistical significance vs. non-diabetic
