## Supplementary Fig. 1 for "Pro-inflammatory cytokines disrupt β-cell circadian clocks in diabetes"

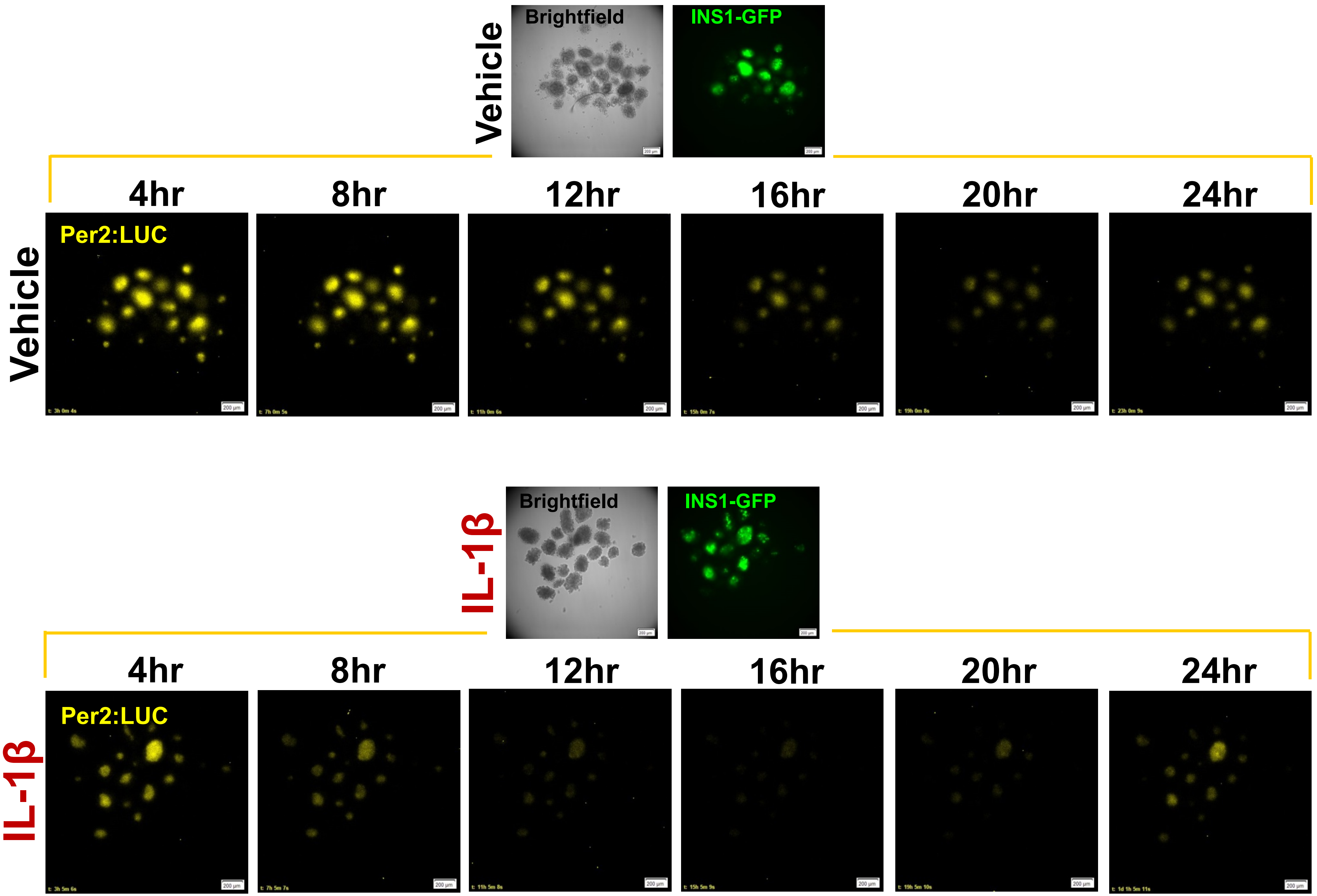


**Supplementary Figure 1.** Representative examples of 24 hours of time-lapse microscopy recordings of circadian *Per2*-driven bioluminescence rhythms in batches of 10-15 islets isolated from *Per2*:LUC-MIP:GFP mice exposed to either vehicle (top) or treated with IL-1β (2 ng/ml) (bottom).
