## Supplementary Fig. 2 for "Pro-inflammatory cytokines disrupt β-cell circadian clocks in diabetes"

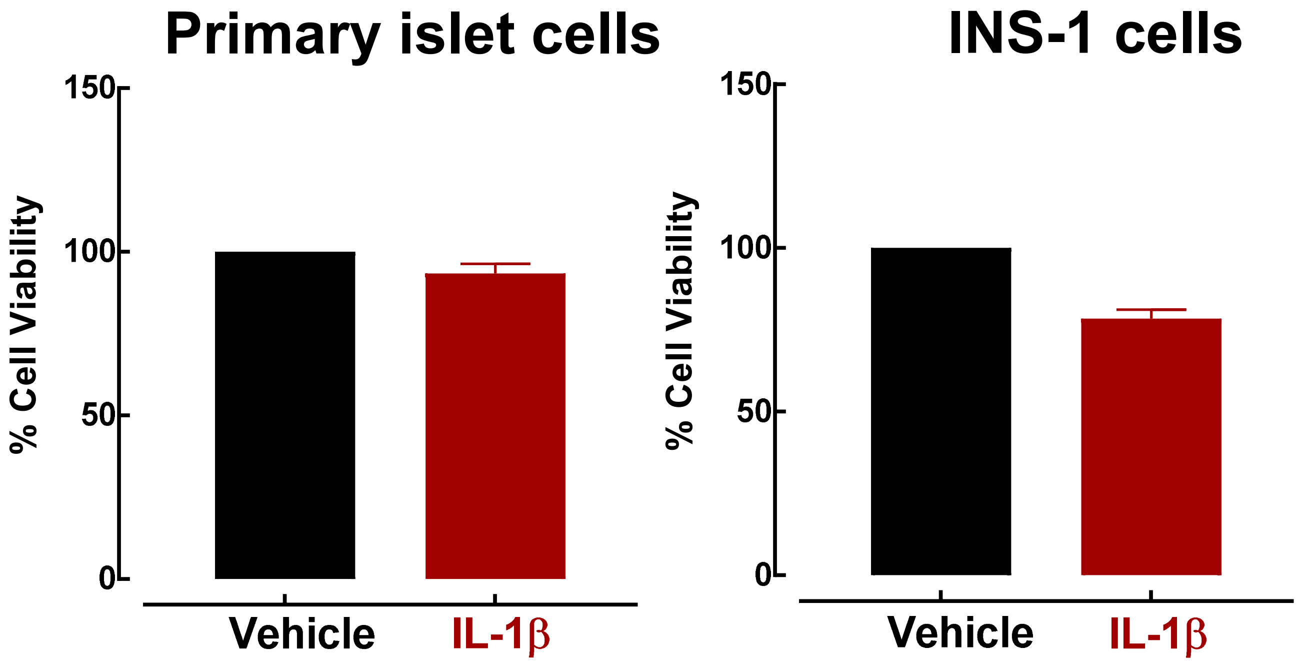


**Supplementary Figure 2. Effects of IL-1β treatment on islet cell viability.** MTT Cell Viability Assay (Cayman Chemicals, Ann Arbor, MI) was performed on dispersed isolated mouse islets (left) and INS-1 832/13 β-cells treated with or without IL-1β (0.2 ng/ml) for 24 h. Primary islets were dissociated with TrypLE solution (Life Technologies) and subsequently plated in triplicate for each condition in a 96 well plate. The protocol for MTT assay was followed as per manufacturer instructions.
