## Supplementary Fig. 3 for "Pro-inflammatory cytokines disrupt β-cell circadian clocks in diabetes"

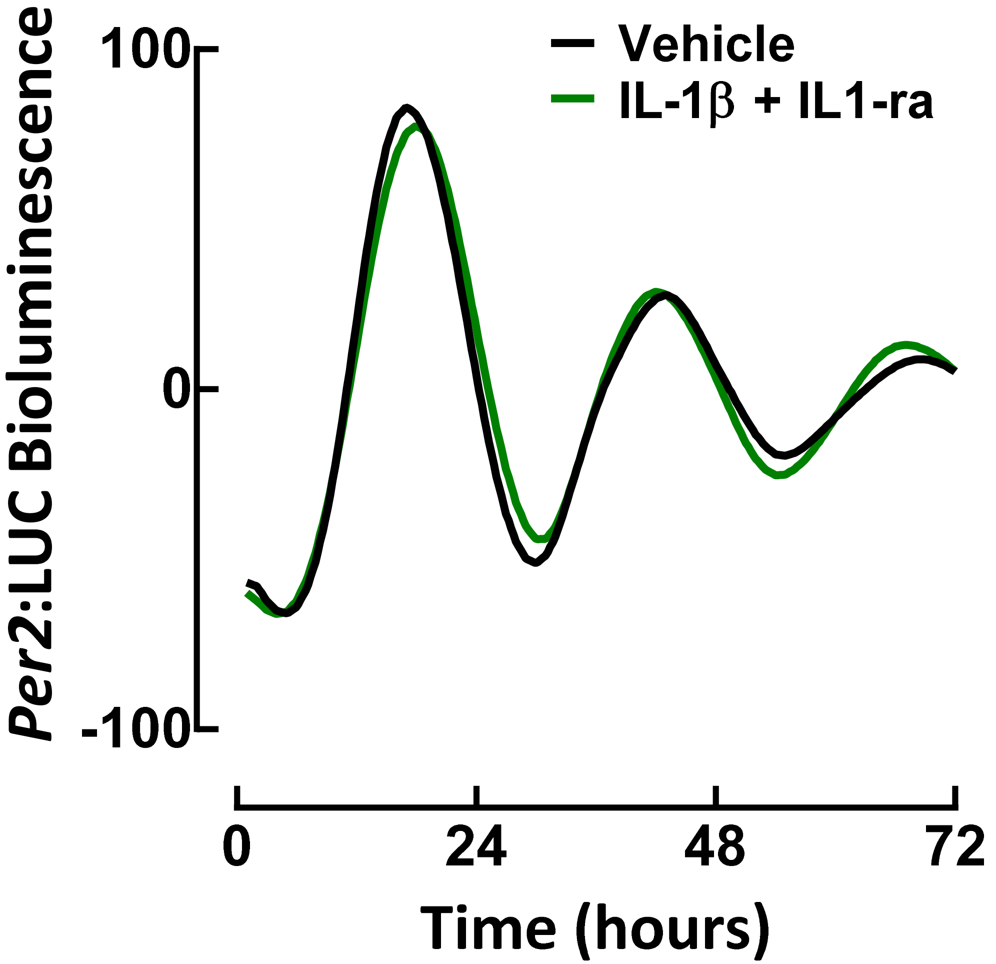


**Supplementary Figure 3. Pretreatment with a specific IL-1 receptor antagonist (IL-1ra) prevents IL-1β-mediated decline in *Per2*-driven bioluminescence in β-cells.** A representative example of circadian *Per2*-driven bioluminescence rhythms in islets isolated from *Per2*:LUC-MIP:GFP mice exposed for 72 hours to either IL-1β (2 ng/ml) + IL-1 receptor antagonist (IL1-ra: 5μg/ml) versus vehicle. Note clear absence of amplitude reduction and phase advance of *Per2*:LUC expression in islets exposed to IL-1β + IL1-ra.
