## Supplementary Fig. 4 for "Pro-inflammatory cytokines disrupt β-cell circadian clocks in diabetes"

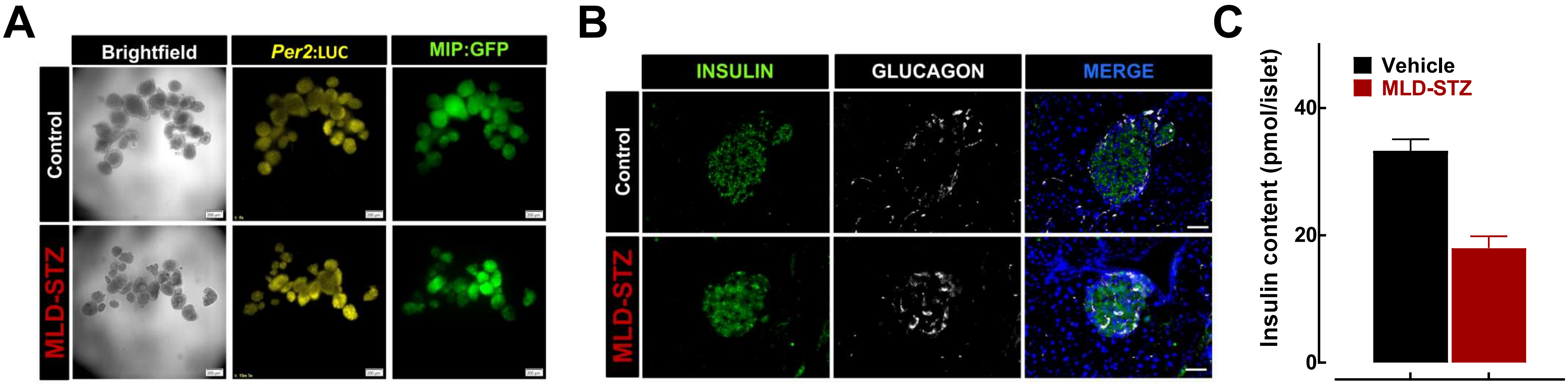


**Supplementary Figure 4. Effects of multiple low dose streptozotocin (MLD-STZ) on islet cell integrity and islet insulin content.** (**A**) Representative images of primary mouse islets used to generate bioluminescence data in Figure 2. Pancreatic islets were isolated from *Per2*:LUC-MIP:GFP mice exposed *in vivo* to MLD-STZ (red) versus vehicle controls. Note preservation of islet integrity and MIP:GFP signal in MLD-STZ islets. (**B**) Representative examples of paraffin embedded pancreatic sections obtained from C57B6 mice exposed *in vivo* to MLD-STZ (red) versus vehicle controls 14 days following STZ treatment. Sections were stained by immunofluorescence for insulin (green), glucagon (grey) and nuclear marker DAPI (blue). Note preservation of islet integrity and attenuated, but preserved insulin signal in MLD-STZ islets. (C) Insulin content measured from 4 independent batches (10-15 islets each) of islets isolated from C57B6 mice exposed *in vivo* to MLD-STZ (red) versus vehicle controls. Insulin content was measured by mouse insulin ELISA (ALPCO diagnostics).
