## Supplementary Fig. 5 for "Pro-inflammatory cytokines disrupt β-cell circadian clocks in diabetes"

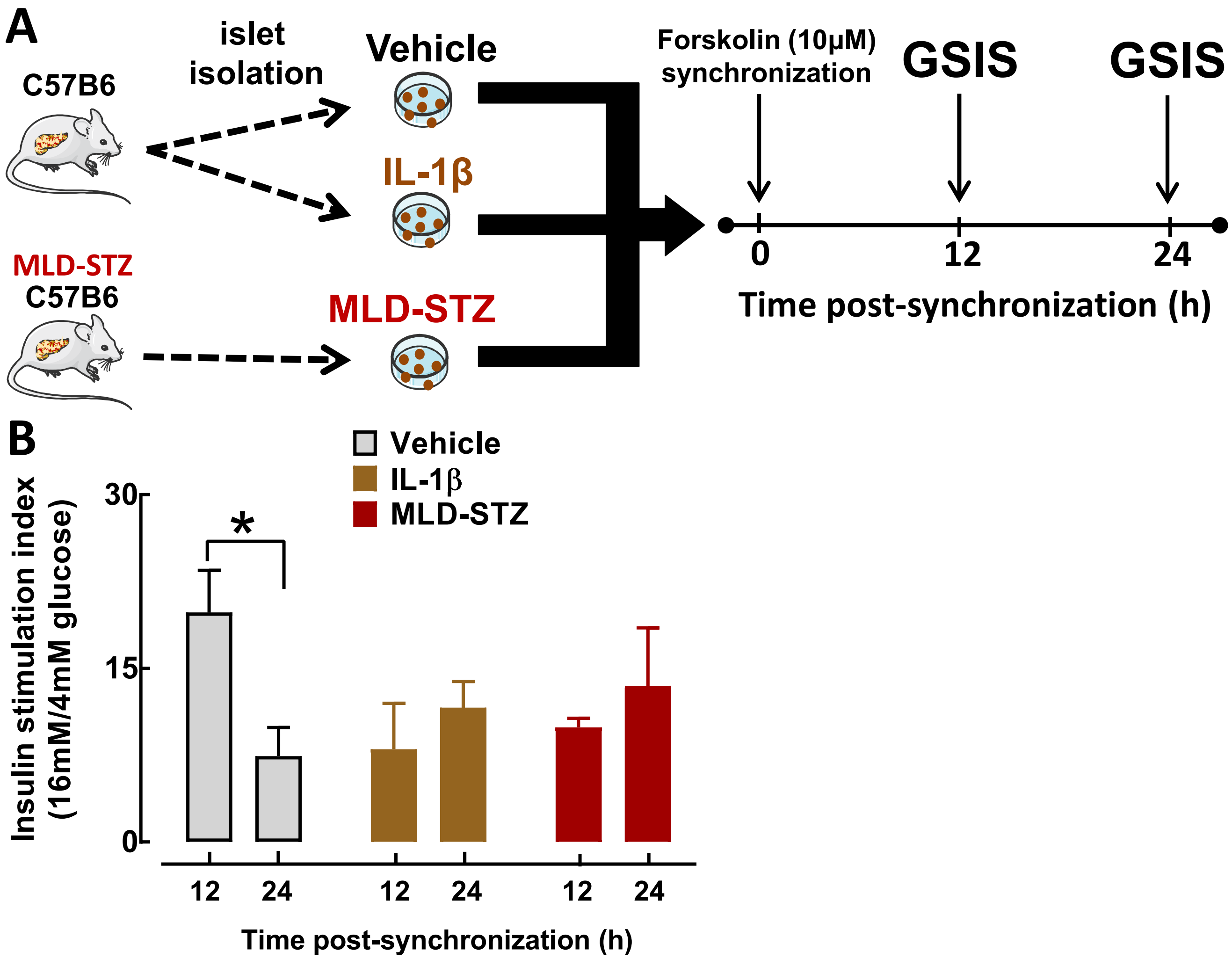


**Supplementary Figure 5. Effects of IL-1β and multiple low dose streptozotocin (MLD-STZ) treatment on circadian regulation of glucose-stimulated insulin secretion (GSIS) in isolated mouse islets**. (**A**) Diagrammatic representation of the study design indicating that isolated islets from control, IL-1β (24 hr treatment in vitro at 2 ng/ml) and MLD-STZ (i.p. STZ; 40 mg/kg/day for 5 consecutive days) treated mice were synchronized with forskolin (10 μM) and glucose-stimulated insulin secretion (GSIS) assessed at 12 and 24 hours post synchronization representative of the peak and trough of islet GSIS in vitro post forskolin synchronization (Perelis et al, 2015). (**B**) GSIS index expressed as insulin release (as % insulin content) during 30 min incubations at hyperglycemic 16 mM glucose over basal 4 mM glucose concentrations assessed at 12 and 24 hours post forskolin synchronization in Control (gray), IL-1β (brown) and MLD-STZ (red) mouse islets. Data are expressed as mean ± SEM and an average of 3 independent experiments (from 3-4 mice per group). $^{*}P < .05$ vs. 24 h post synchronization.
