## Supplementary Fig. 6 for "Pro-inflammatory cytokines disrupt β-cell circadian clocks in diabetes"

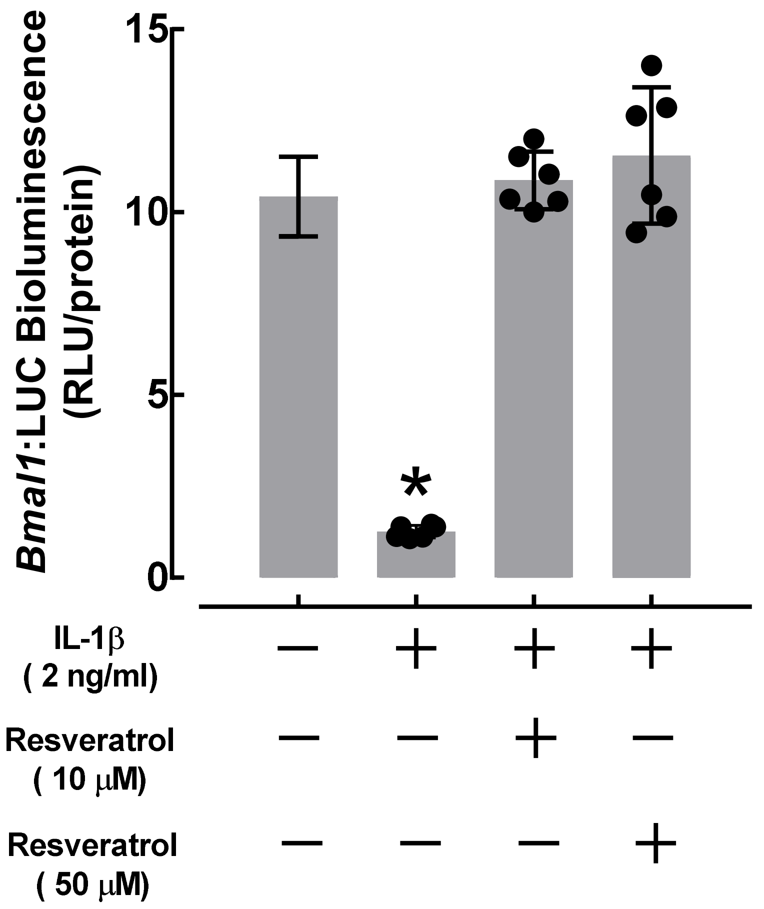


**Supplementary Figure 6. Sirt1 activator Resveratrol reverses deleterious effects of IL-1β on Bmal1 promoter activation.** Bmal1 promoter activity assessed using stable-transfected Bmal1:luciferase (Bmal1:LUC) reporter expressing INS-832/13 β-cells exposed for 24 h to IL-1β (0.1-5 ng/ml) with or without co-treatment with Sirt1 chemical agonist Resveratrol (10 and 50μM). Values are mean ± SEM ($n$=6 independent experiments per given condition) and *$P$<0.05 denotes statistical significance vs. untreated.
