## Supplementary Fig. 7 for "Pro-inflammatory cytokines disrupt β-cell circadian clocks in diabetes"

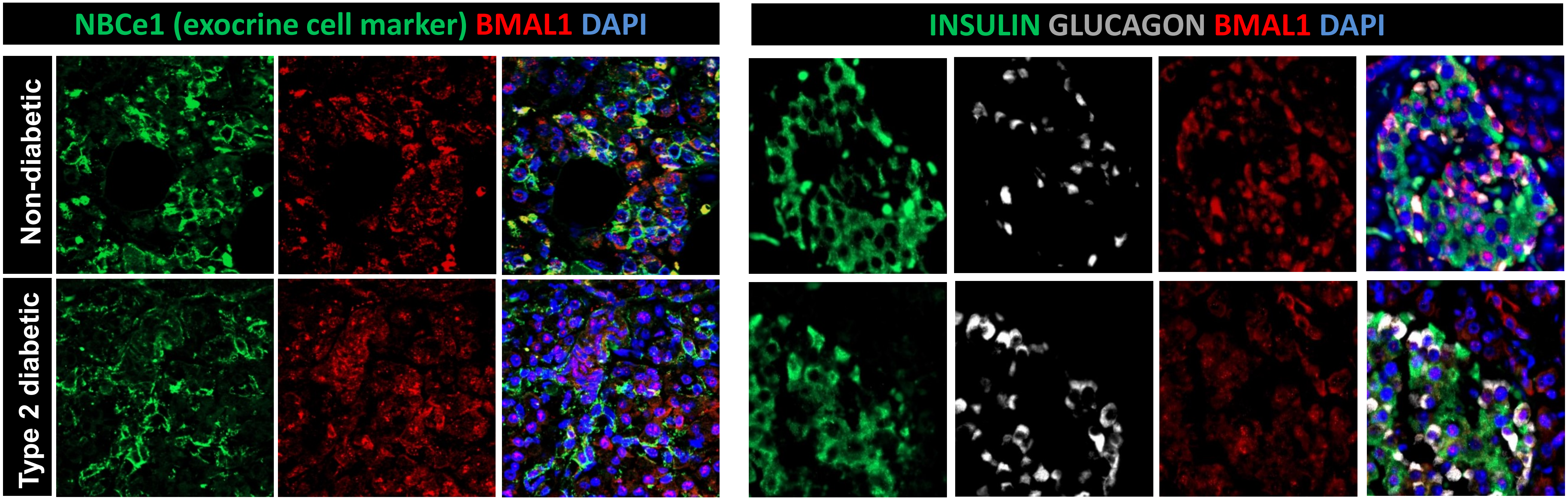


**Supplementary Figure 7. Preserved BMAL1 expression in the exocrine pancreas of patients with Type 2 diabetes.** (**Left panel**) Representative examples of paraffin embedded human pancreatic sections from nondiabetic subjects and subjects with type 2 diabetes stained by immunofluorescence for pancreatic exocrine cell marker NBCe1 (green), BMAL1 (red) and nuclear marker DAPI (blue). Note comparable expression of BMAL1 in the exocrine pancreas of nondiabetic and type 2 diabetic subjects. (**Right panel**) Representative examples of paraffin embedded human pancreatic sections from nondiabetic subjects and subjects with type 2 diabetes stained by immunofluorescence for Insulin (green), Glucagon (grey), BMAL1 (red) and nuclear marker DAPI (blue).
