## Supplementary Fig. 8 for "Pro-inflammatory cytokines disrupt β-cell circadian clocks in diabetes"

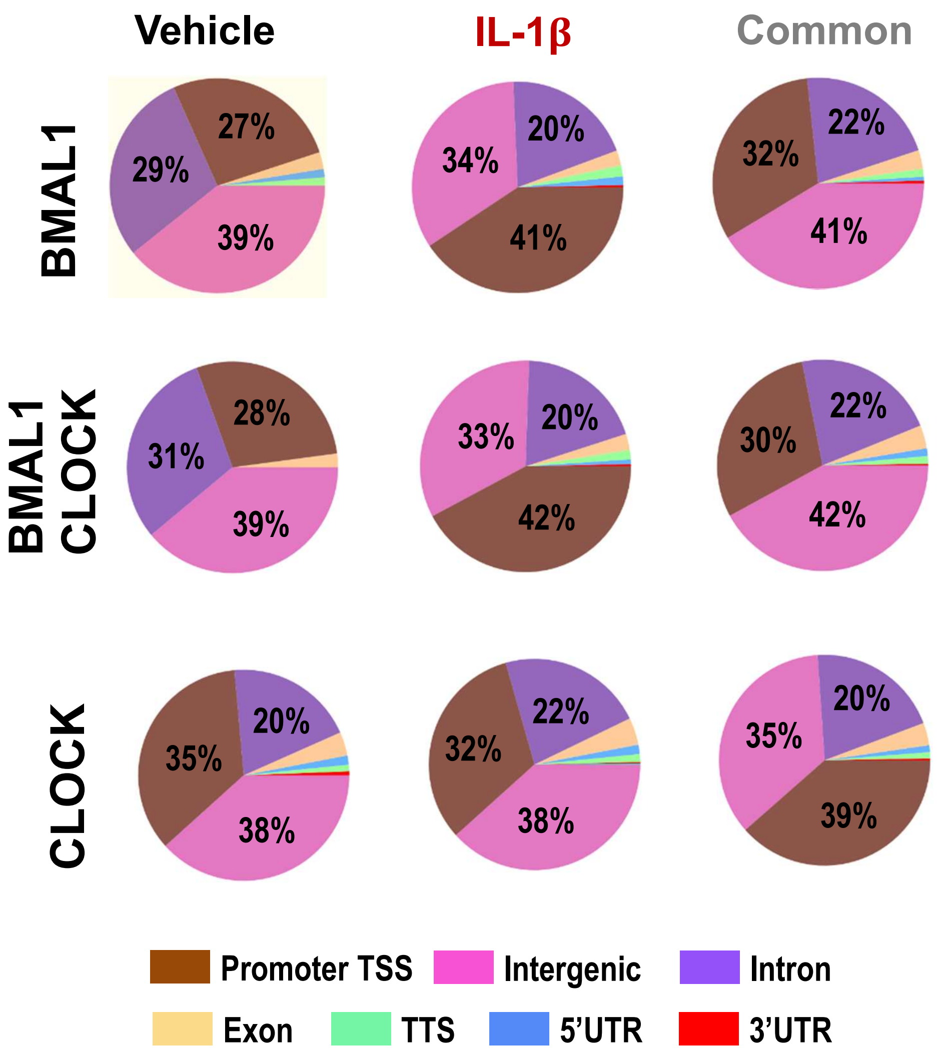


**Supplementary Figure 8. Genome-wide distribution of Bmal1/CLOCK binding peaks.** Global genome-wide distribution of BMAL1, CLOCK and BMAL1/CLOCK cobound binding peaks throughout the genome under vehicle and/or IL-1β conditions.
